# Transient overdominance, coadaptation, and the fixability of heterosis

**DOI:** 10.1101/2023.08.23.554444

**Authors:** Hilde Schneemann, John J. Welch

## Abstract

Many species pairs form F1 hybrids that are fitter than their parents. Such heterosis can arise if the parents carry recessive deleterious mutations; and in this case, the heterosis should be fixable, because selecting out the deleterious mutations yields a high-fitness homozygous hybrid. However, heterosis might not be fixable if caused by overdominance (an intrinisic advantage to heterozygosity) or if the parents contain coadapted gene complexes. These alternatives have been tested with introgression lines, where small regions of genome are scored in the heterospecific background. We develop predictions for introgression line data under a simple model of phenotypic selection, where parents diverge by fixing deleterious mutations via genetic drift. We show that this simple process can generate complex patterns in the data, misleading tests for both overdominance and coadaptation. We also suggest new ways to analyse the data to overcome these difficulties. Reanalyses of published data from *Solanum* and *Gossypium* suggest that the model can account for the qualitative patterns observed, though not the extent of apparent overdominance.

## Introduction

Many divergent populations or closely-related species can form first-generation hybrids, and sometimes these F1 are fitter than their parents (Darwin, 1876; Gowen, 1952; Lippman and Zamir, 2007; Mackay et al., 2020; see Figure 1A). This F1 heterosis can have important implications for the outcomes of wild hybridization (Thompson and Schluter, 2022; Kulmuni et al., 2023), and is exploited by plant and animal breeders (East, 1908; Shull, 1908; Gowen, 1952; Gerdes et al., 1999; Gur and Zamir, 2004; Mackay et al., 2020), and by conservationists (Genovart, 2008; Chan et al., 2019). Because F1 are fully heterozygous, they cannot breed true, and so an important question is whether the high fitness might persist into later hybrid generations. We say that heterosis is “fixable” if there exists a hybrid, which is homozygous at all loci and shares the high fitness of the F1, and if this fit homozygous hybrid can be bred from the F1 generation (Birchler, 2003; Koltunow and Tucker, 2003; Mackay et al., 2020; Figure 1A).

**Figure 1:**
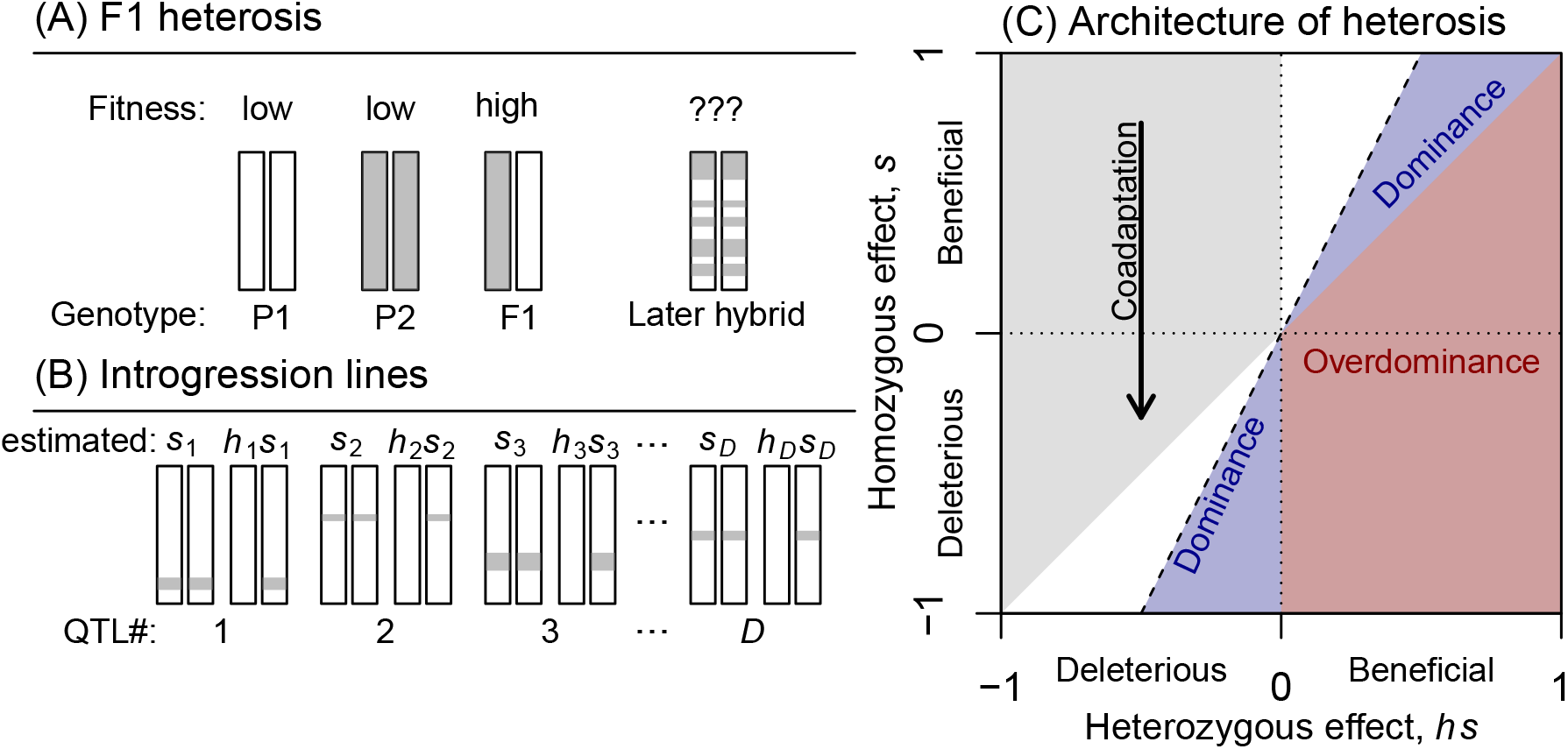
Introgression lines can be used to investigate the causes of F1 heterosis. **(A)** First generation, F1 hybrids between parental lines P1 and P2 can show heterosis, or higher fitness. The heterosis is “fixable” if we could evolve a later-generation, homozygous hybrid with similarly high fitness. **(B)** Introgression lines can be used to score small regions of genome from a donor line (P2) introgressed into a recipient line (P1), in both heterozygous and homozygous state. **(C)** Plotting the homozygous effects of these QTL against their heterozygous effects can be used to test theories of heterosis. QTL falling in the red region support the “overdominance theory” of heterosis, in which heterozygosity is intrinsically beneficial. QTL falling into the two blue triangles support the “dominance theory” of heterosis in which one or other of the parental lines has fixed recessive deleterious mutations. Coadaptation between the parental alleles, which makes heterosis difficult to fix, can be inferred if most QTL fall in the lower half of the plot (see arrow), even when the parental lines having similar fitness.

Whether heterosis is fixable depends on how it is caused (Crow, 1948; Koltunow and Tucker, 2003; Lippman and Zamir, 2007), which may also depend on how the parental lines diverged (Crow, 1948; De Sanctis et al., 2023). Under the classical “dominance theory” of heterosis, the high F1 fitness is explained by the masking of recessive deleterious mutations, when present as heterozygotes (Bruce, 1910; Keeble and Pellew, 1910; Crow, 1948). This theory therefore implies that heterosis will be substantial only when the parents have fixed recessive deleterious mutations via genetic drift (Crow, 1948). The theory also implies that the heterosis should be fixable, because the deleterious mutations could be selected out, leaving a high-fitness homozygous hybrid, containing only the ancestral alleles. By contrast, under the “overdominance theory” of heterosis, heterozygous genotypes enjoy an intrinsic fitness advantage over both homozygotes (East, 1908; Shull, 1908; Crow, 1948). This theory makes no assumptions about the divergence process, but it does imply that the heterosis will not be fixable, because later generation hybrids will lose the beneficial heterozygosity.

However the heterosis is caused, it might also be difficult to fix if the parental genotypes contain coadapted gene complexes – i.e. if there was past selection for sets of alleles that function well together, but poorly in novel combinations (Hill, 1982; Lynch, 1991; Simon et al., 2018). Negative epistasis between heterospecific alleles creates segregation load after hybridization, and a severe loss of fitness for almost all post-F1 hybrids (Slatkin and Lande, 1994; Chevin et al., 2014; Simon et al., 2018; Schneemann et al., 2020).

To understand the fixability of heterosis, the alternatives above have been tested in several ways (Hill, 1982; Birchler, 2003; Lippman and Zamir, 2007; Wang et al., 2012; Jiang et al., 2017; Torgeman and Zamir, 2023), but one powerful approach uses introgression lines, where small regions of genome from a donor parental line are introgressed into a recipient parental line, e.g. using repeated backcrossing (Eshed and Zamir, 1994; Semel et al., 2006; Lippman and Zamir, 2007; Wang et al., 2012; Alseekh et al., 2013; Tian et al., 2019; Figure 1B). The fitness of each QTL is then scored in both homozygous and heterozygous state (Figure 1B). The architecture of heterosis is revealed by plotting the homozygous effects, *s*, against the heterozygous effects, *hs* (Figure 1C). All QTL falling to the right of the dashed line in Figure 1C will act to create F1 heterosis. Of these, QTL in the red area of the plot are overdominant, and so support the overdominance theory of heterosis. By contrast, QTL in the two blue areas support the dominance theory of heterosis. While the dominance theory posits only a single class of mutation – deleterious recessives – there are two distinct areas because introgressions could involve either derived alleles (when the deleterious recessive fixed in the donor line), or ancestral alleles (when the deleterious recessive fixed in the recipient line). If the dominance theory holds, then only compound QTL, containing both ancestral and derived alleles, could fall in the red area, and so manifest associative-or pseudo-overdominance (Jones, 1917; Collins, 1921; Lippman and Zamir, 2007).

Introgression lines, even when they contain single QTL, can also be used to test for coadaptation between the parental alleles. In particular, if parental alleles are coadapted, then we should expect most homozygous introgressions to be deleterious, even when the donor and recipient lines have similar levels of fitness. As such, coadaptation can be inferred if QTL tend to fall preferentially in the lower half of Figure 1C.

In the rest of this paper, we compare the simple and intuitive predictions above to the explicit predictions of a mathematical model of selection. In particular, we will use a well-known and well-studied model of selection acting on quantitative traits (Fisher, 1930; Lande, 1980; Turelli, 1985; Slatkin and Lande, 1994; Orr, 1998; Barton, 2001), which makes similar predictions to other phenotypic models (Martin, 2014; Cotto and Day, 2023). We will focus on cases where the parental lines diverge under the action of genetic drift – consistent with the dominance theory (Crow, 1948). Previous work has shown that this model can generate heterosis, with or without coadaptation (Barton, 2001; Chevin et al., 2014; Simon et al., 2018; Schneemann et al., 2022; De Sanctis et al., 2023), and that it can generate transient overdominance – i.e. overdominance that comes and goes with changes in the genetic background (Manna et al., 2011; Sellis et al., 2011). It therefore has the potential to mislead tests of the theories of heterosis using introgression lines. Our goal is to ask how far the tests could be misled in practice.

## Model and Results

### Definition of heterosis and composite effects

In the following sections, we will compare predictions for introgression lines to predictions for the composite effects, as defined by Hill (1982). These effects quantify the interactions between heterospecific alleles in arbitrary hybrid genotypes, and resemble the variance components of standard quantitative genetics (Lynch, 1991; Lynch and Walsh, 1998). The expected log fitness of any class of hybrid can be written as a weighted sum of the composite effects (Hill, 1982; Schneemann et al., 2020), and so calculating the composite effects can tell us a lot about the causes of F1 heterosis and about its fixability.

We will consider the composite effects as they apply to log fitness – because fitness differences are best measured on a multiplicative scale (Phillips et al., 2000), and we restrict ourselves to the first- and second-order effects (involving interactions between no more than two loci); this is often a good approximation, and is exact for the simple phenotypic model we will introduce below. In particular, let us consider a balanced (50:50) hybrid, in which a fraction 0 *≤ π ≤* 1 of the loci are heterozygous for heterospecific alleles. If we denote the fitness of this hybrid as *w*^*π*^, and the fitness of the original parental lines as *w*^P1^ and *w*^P2^, then we have

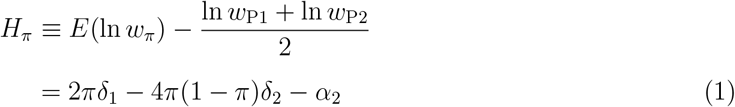

where *δ*^1^ is the dominance composite effect, *δ*^2^ is the dominance-by-dominance interaction, and *α*^2^ is the additive-by-additive epistatic interaction (Hill, 1982; Schneemann et al., 2020; De Sanctis et al., 2023). Since the F1 is fully heterozygous, we have *π* = 1 and its midparent heterosis is simply

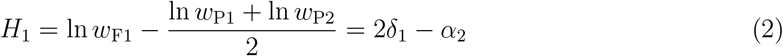

while the equivalent quantity for the second-generation F2 is *H*_0.5_ = *δ*_1_*−δ*_2_*−α*_2_. Finally, *H*_0_ = *−α*_2_, describes the expected log fitness difference between the parents and an entirely homozygous genotype, assembled from a random mix of their alleles; *α*_2_ is therefore a natural measure of the coadaptation among the parental alleles (Lynch, 1991). Large positive values of *α*_2_ not only reduce F1 heterosis (eq. 2), but also create segregation load among later generation hybrids (Slatkin and Lande, 1994; Chevin et al., 2014).

### The phenotypic model

To assign fitnesses to genotypes, we will follow many previous authors (e.g. Lande, 1980; Turelli, 1985; Orr, 1998) and use a model of quadratic selection acting on *n* quantitative traits. Under this model, an individual’s log fitness is given by

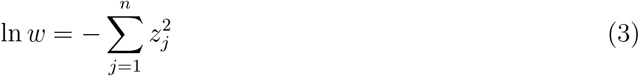

where *z*_*j*_ is the deviation of trait *i* from its optimal value. We will assume that each introgression from the donor to the recipient line has an additive and a dominance effect on each trait, such that, for the *i*^*th*^ such introgression, trait *j* is altered by a value 2*a*_*ij*_ when the introgression is homozygous, and a value *a*_*ij*_ + *d*_*ij*_ when the introgression is heterozygous. The composite effects of eq. 1 are defined as various summary statistics of the *a*_*ij*_ and *d*_*ij*_ (see De Sanctis et al., 2023 and Appendix eqs. 27-29).

The values that actually characterise two divergent populations will depend on the mode of divergence between them; to see this, let us now consider the predictions of the model under two explicit models of divergence.

### Predictions under random mutation accumulation

Let us first consider results under eq. 3 when parental lines diverged via random mutation accumulation, as might occur if population size were very small, or inbreeding severe (Oakley et al., 2015, 2019). In particular, we assume that the parental lines began with an optimal phenotype, and then accrued *D* randomly orientated mutations. We further assume that the additive effect of these mutations, the *a*_*ij*_, are normally distributed with a mean of zero, and a variance of *ν*. This implies that the mean selective effect of a new mutation, if placed in homozygous state in an optimal background, is

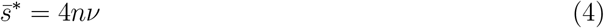

where the asterisk is used to distinguish these selective effects from the selective effects of the introgressions. We will now assume that the phenotypic dominance effects, the *d*_*ij*_, of each mutation are also independently normally distributed, with zero mean and variance *νβ*. The parameter *β* therefore describes the relative sizes of the additive and dominance phenotypic effects. With these assumptions, it follows from previous results (De Sanctis et al., 2023; Schneemann et al., 2022; see also Appendix for details), that the expected composite effects are:

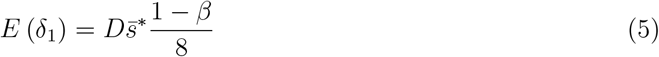

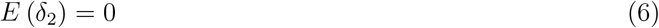

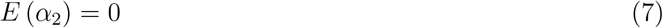

And so, from eq. 2, the expected F1 heterosis is

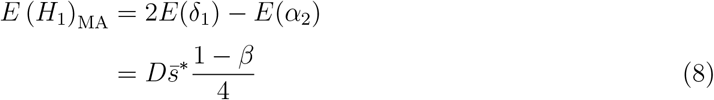

(where the subscripted MA indicates random Mutation Accumulation). Equation 8 shows that F1 heterosis is expected if the phenotypic dominance effects are typically smaller than the additive effects (i.e. if *β <* 1). The heterosis captured in eq. 8 is also likely to be fixable. In particular, while epistasis can exist between pairs of mutations, the epistasis will fluctuate in sign, and it follows from eq. 7, there is no coadaptation on average (Martin et al., 2007; Simon et al., 2018; Fraïsse and Welch, 2019; De Sanctis et al., 2023). As such, there will be no additional load for post-F1 hybrids beyond that caused by the loss of heterozygosity (Simon et al., 2018; De Sanctis et al., 2023). Populations are therefore likely to fix the ancestral allele at each locus, recovering a high fitness homozygous hybrid.

The comments above hold for all parameter values, but the predicted architecture of heterosis (Figure 1C) varies qualitatively in different parameter regimes. To see this, Figure 2A-D summarizes results that are derived in the Appendix. Figure 2A shows an illustrative simulation where *D* (the number of mutations that differentiate the lines) is much smaller than *n* (the number of phenotypic traits under selection). When *D/n ≪* 1, the distribution of single QTL (i.e. introgressions containing only a single divergent site) comprises two distinct clusters in the two blue areas, corresponding to the introgression of either derived or ancestral alleles. In this parameter regime, introgressions of derived alleles are always deleterious 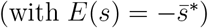 and the introgressions of ancestral alleles beneficial 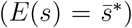. The dominance coefficients also differ for the deleterious and beneficial introgressions. To see this, consider a simple estimator of the typical dominance coefficient

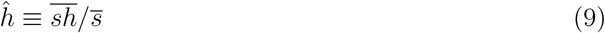

where we have used a simple ratio of the sample means of the heterozygous and homozygous effects (see, e.g., Manna et al., 2011 eq. 2). If we now restrict the estimates to beneficial or deleterious introgressions, then we find

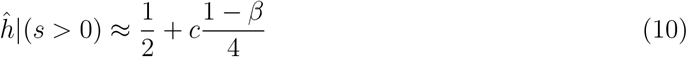

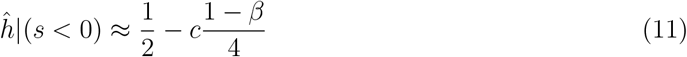

where *c >* 0 is a constant which varies with parameter regime. We show in the Appendix that *c* = 1 holds when *D < n*, and introgressions comprise single alleles. It follows, therefore, that when heterosis is expected (i.e., when *β <* 1; eq. 8) then introgressions of ancestral alleles are dominant, while introgressions of derived alleles are recessive.

**Figure 2:**
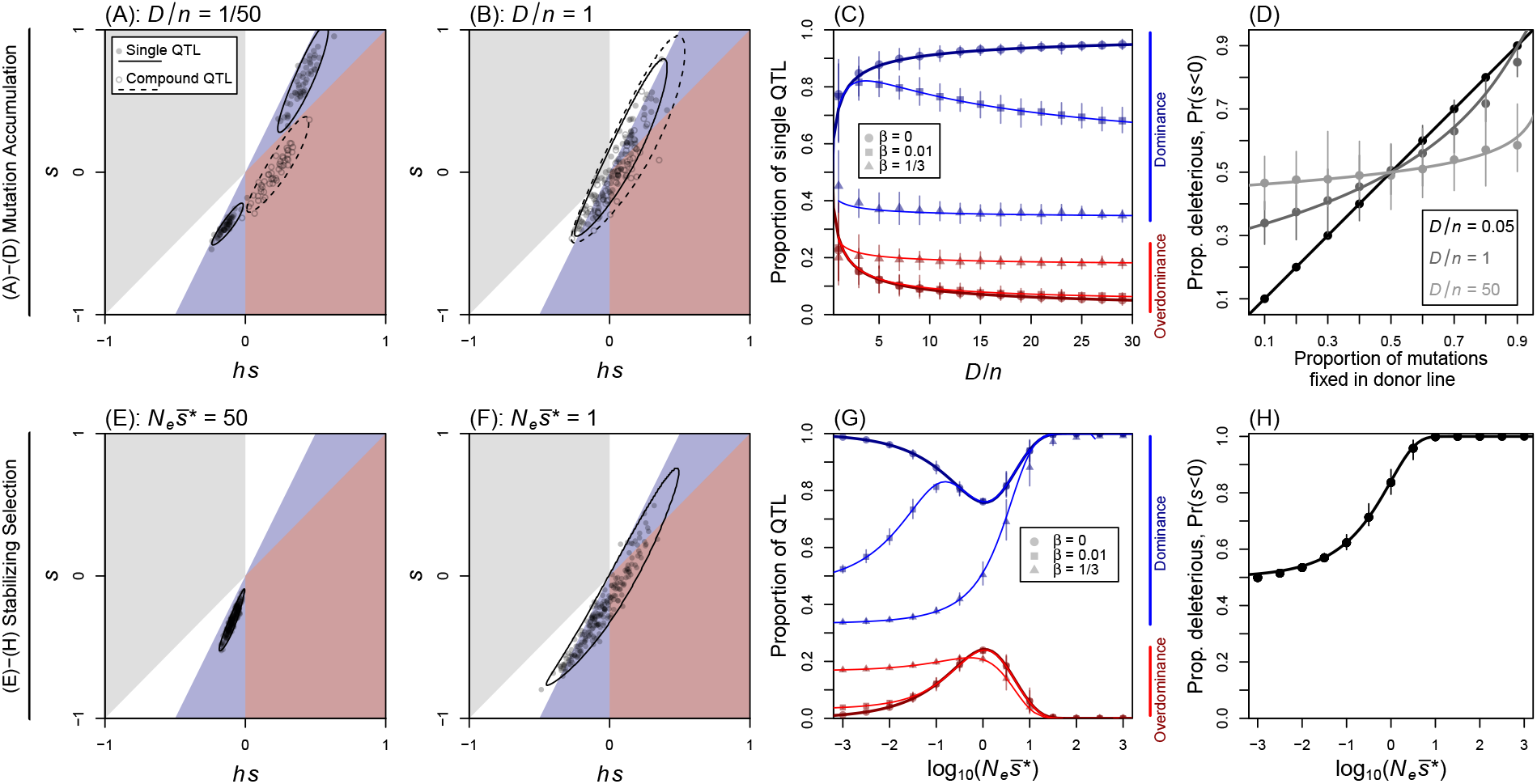
The architecture of heterosis under a simple model of phenotypic selection. Predictions for introgression lines derived from parents that diverged by **(A)-(D)**: random mutation accumulation, or **(E)-(H)**: partially effective stabilizing selection. Analytical predictions (lines) are compared to simulations (points); with error bars indicating the mean and range over 50 replicates. **(A)** Under random mutation accumulation, when selection acts on many traits (*D* = 100 *≪ n* = 5000), results agree with predictions of the classical dominance theory of heterosis, with QTL containing single derived or ancestral alleles falling in the two blue areas, and compound QTL (containing one derived and one ancestral allele) showing pseudo-overdominance. **(B)** But with fewer traits (*D* = *n* = 100), the architecture becomes complex, with both single and compound QTL found in all areas of the plot. **(C)** The proportions of QTL supporting the dominance and overdominance theories depend solely on the ratio *D/n* and the relative size of the dominance effects, *β*. **(D)** The proportion of homozygous introgressions that are deleterious, accurately reflects the proportion that fixed in donor population when *D < n*, because derived mutations tend to be deleterious; however, when *D ≫ n* the proportion approaches 1*/*2 regardless. **(E)** Under stabilizing selection, if 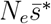 is large (i.e. if introgressions would be under effective selection in an optimal background), then all introgressions are deleterious and recessive, indicative of coadaptation between the parental alleles. **(F)** But when 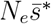 ≈ 1, the architecture is also complex. **(G)** Support for the dominance and overdominance theories depends solely on 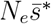, and *β*. **(H)** The proportion of introgressions that are deleterious increases with 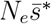, so that coadaptation is difficult to detect when 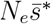 < 1. (Other parameters, chosen for visualisation, are (A): *β* = 1*/*5, *ν* = 0.12*/n*; (B): *β* = 1*/*5, *ν* = 0.01*/n*; (C): *n* = 50; (D): *D* = 100; (E): *D* = 200, *n* = 20, *ν* = 0.1*/n, β* = 0.0005; (F): as (E) but with *β* = 0.025; (G)-(H): *D* = 20000, *n* = 40, *N* = 10 and *N* = 100. See Appendix for full details).

When introgressions contain both ancestral and derived alleles, then the same model generates pseudo-overdominance (Jones, 1917; Collins, 1921). To see this, the empty points in Figure 2A show the same set of simulated introgressions, but now combined into compound QTL containing exactly one derived allele and one ancestral allele. These compound QTL now fall in the red region of the plot, and this is because, for such QTL, eqs. 10-11 still hold, but with 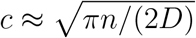 (see Appendix for details). As such, when *D/n <* 0.39(1 *− β*)^2^, *c* will be large enough to produce overdominance for a typical compound QTL. In summary, Figure 2A shows that with random mutation accumulation and *D < n*, results from the phenotypic model agree with the classical dominance theory of heterosis (Bruce, 1910; Keeble and Pellew, 1910; Crow, 1948).

Now let us consider the same scenario, after decreasing the number of traits, such that *D* = *n*. In this case, as shown in Figure 2B, results change qualitatively such that we obtain a complex genetic architecture including both dominant and overdominant QTL. In particular, when *D* = *n*, the distribution of single QTL becomes unimodal, with around 1/5 of the single QTL now yielding apparent support for the overdominance theory. Moreover, results for single QTL become qualitatively similar to results for compound QTL. For example, when *D ≥ n*, then eqs. 10-11 hold for single QTL, but with 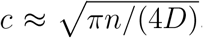. This makes the results for single and compound QTL comparable to one another.

As shown in Figure 2C, these patterns continue as *D/n* increases, and when mutation effects are normal, the proportion of QTL in each area of the plot can be predicted solely from *β* (the relative size of the dominance effects) and the ratio *D/n* (see also Figure S1A). Another difference between the parameter regimes is shown in Figure 2D. When *D < n*, the proportion of QTL that are deleterious is roughly equal to the proportion that fixed in the donor line – because, under random mutation accumulation, derived alleles tend to be deleterious, and ancestral alleles tend to be beneficial. However, when *D ≫ n*, around half of introgressions become deleterious, regardless of where the mutations fixed.

The results in Figure 2B-D arise because, when *D > n*, mutations will often act in similar phenotypic directions, and so show fitness interactions in some genetic backgrounds. Nevertheless, the results do not imply an intrinsic advantage to heterozygosity, but instead represent the transient overdominance that arises naturally with optimizing selection (Manna et al., 2011; Sellis et al., 2011). In recombinant hybrids, derived mutations will be deleterious on average, whether in homozygous or heterozygous state.

### Predictions under partially effective stabilizing selection

Results above apply if populations diverge by fixing random mutations, without the action of natural selection. However, genomic divergence can also occur under stabilizing selection, if genomic changes cause little net phenotypic change (Hartl and Taubes, 1996). Again, the process depends on the action of genetic drift, albeit with compensatory mutations, and is sometimes called “system drift” (Rosas et al., 2010; Schiffman and Ralph, 2021). In the Appendix (see also De Sanctis et al., 2023) we show that eqs. 5 and 6 also hold under system drift, so that the composite effects involving dominance remain unchanged. However, stabilizing selection does create coadaptation among the parental alleles, and we find

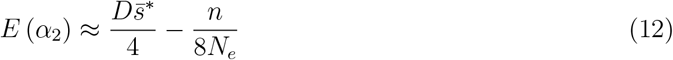

(see also Chevin et al., 2014; Barton, 2016; Simon et al., 2018; De Sanctis et al., 2023). Note that eq. 12 depends on the effective population size, *N*_*e*_, since this determines how well populations track their optima (Barton, 2016; Schneemann et al., 2022; De Sanctis et al., 2023). It now follows from eqs. 2, 5 and 12, that the expected level of F1 heterosis is

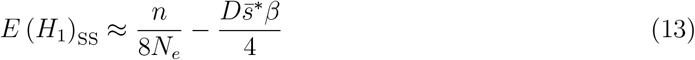

(where the subscripted SS denotes the action of Stabilizing Selection). So heterosis can coexist with coadaptation on the condition that

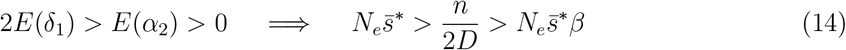

However, as a result of the coadaptation, the heterosis that arises under stabilizing selection need not be fixable in practice. Indeed, if the ancestral genotype is no fitter than the parents, the heterosis might not even be fixable in principle.

Once again, as shown in Figure 2E-H, the architecture of heterosis can vary qualitatively in different parameter regimes. The key parameter under stabilizing selection is 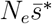, which quantifies the efficacy of selection on the homozygous QTL, and the same results apply to compound QTL, if 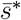 is defined as the selective effect of the whole introgression.

Figure 2E shows an example where 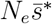 is large, such that selection acts effectively on the QTL. In this case, as with Figure 2A, the data are relatively easy to interpret. In Figure 2E, all QTL fall within a blue region of the plot, accurately reflecting the divergence via (system) drift on recessive mutations. And all introgressions are deleterious, accurately reflecting the strong coadaptation between the alleles carried by the two parents. By contrast, Figure 2F shows results when QTL are nearly neutral 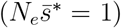, and there the architecture is complex, and qualitatively similar to that shown in Figure 2B. Figure 2G (see also Figure S1B-D) shows that complex architectures can also appear when 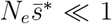, but only when *β* is large, implying that heterosis is less likely to appear in this parameter regime (eq. 14). Figure 2H shows that the evidence for coadaptation also depends on 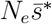, and that it disappears when 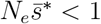.

### Detecting coadaptation from introgression lines

The preceding sections show that similar genetic architectures of heterosis can arise, despite very different levels of coadaptation between the parental alleles (Figures 2B and 2F). When architectures are complex, the proportion of QTL that are deleterious is a poor indicator of coadaptation, because we can observe Pr(*s <* 0) *≈* 1*/*2 even when coadaptation is strong (Figure 2H).

Nevertheless, introgression line data can be used to infer coadaptation in a more direct way. Most straightforwardly, if parental and F1 fitnesses are available, we can compare the observed level of F1 heterosis to the level predicted from the individual QTL, assuming no fitness interactions between them. In particular, we show in the Appendix that, if there are no fitness interactions between loci, then we have

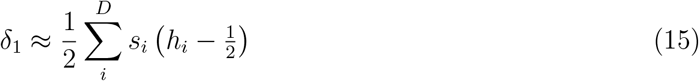

As such,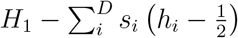 can be used as an estimator of *α*_2_ (eq. 2).

A second, even simpler approach exploits the correlations between dominance and epistatic effects under phenotypic models, and in particular, the relation that 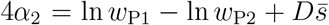 (Hill, 1982; Chevin et al., 2014; De Sanctis et al., 2023; see also Appendix). It follows that, when the parents have similar fitness, such that *w*_P1_ *≈ w*_P2_, then the level of coadaptation can be estimated solely from the mean dominance coefficient. To see this, let us use the estimator of the dominance coefficient of eq. 9, and define the quantity, *r*:

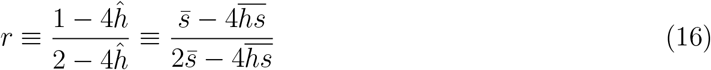

which we show in the Appendix is an estimator of the following quantity

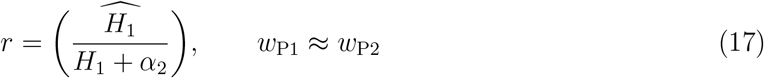

It follows that if we observe heterosis, with a complex genetic architecture, then the presence of substantial coadaptation (i.e. negative epistasis between the heterospecific alleles) should lead to 0 *< r ≪* 1, while the absence of coadaptation can be inferred if *r ≈* 1. This is confirmed by simulations reported in Figure S2. Unlike our predictions for the proportions of QTL that fall in different regions of the graph (Figures 2 and S1), Figure S2 shows that eq. 17 is quite robust to deviations from our assumption of normality in the substitution effects.

### Illustrative data reanalysis

In this section and Figure 3, we consider two published data sets from introgression lines (Semel et al., 2006; Tian et al., 2019). These data present, arguably, the strongest evidence for multilocus overdominance contributing to heterosis (e.g., Kaeppler, 2012). Our aim is to show how the theoretical results above might affect our interpretation of such data.

**Figure 3:**
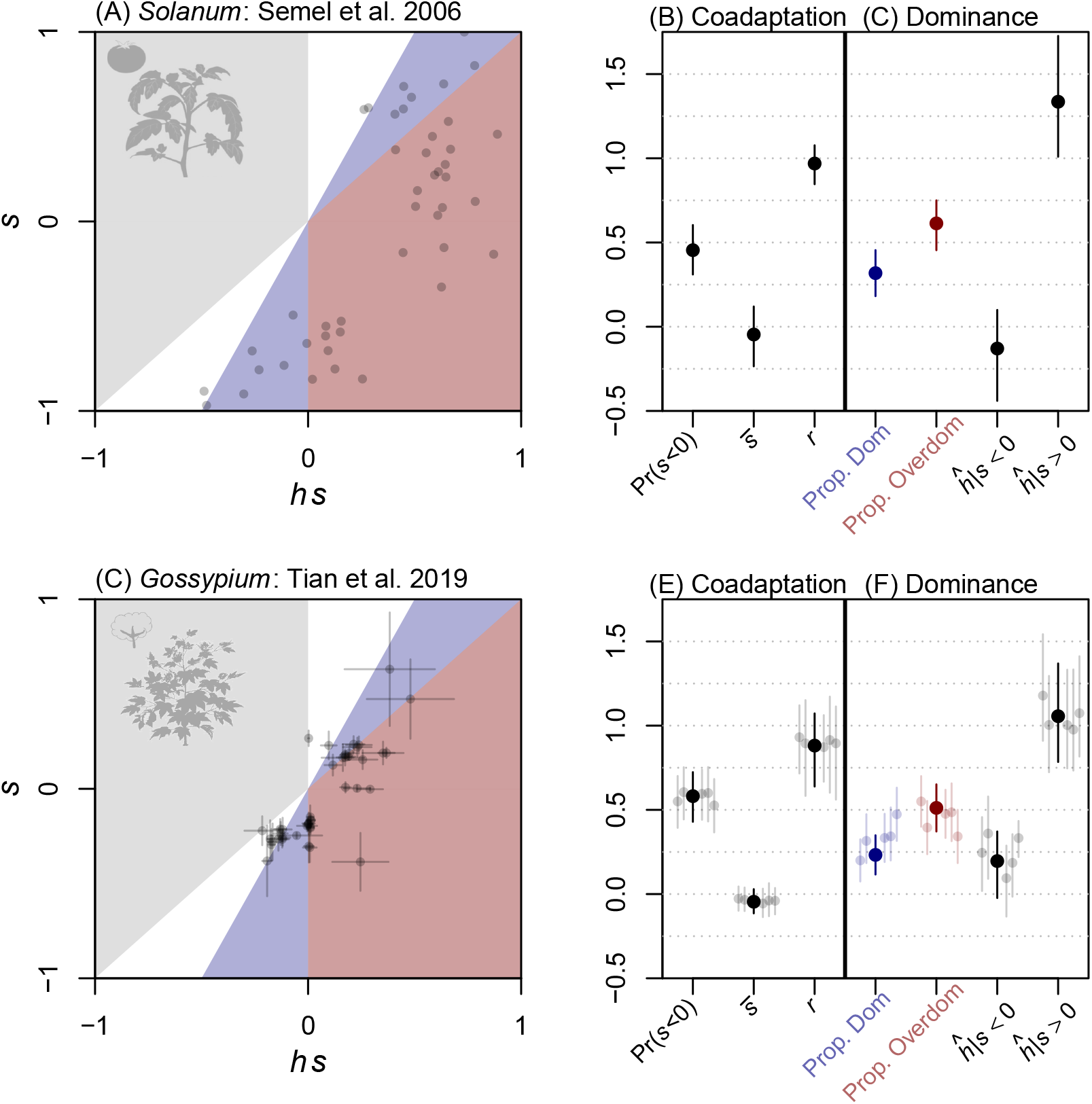
Empirical architectures of heterosis. **(A)-(C)**: Published data from 44 introgression lines, involving small genomic regions from the wild tomato *Solanum pennellii* introgressed into the cultivated tomato *Solanum lycopersicum* (Semel et al., 2006); seed weight per fruit is the fitness proxy. **(D)**-**(F)**: Published data from 43 introgression lines, involving small genomic regions from the New World cotton species *Gossypium barbadense* introgressed into *Gossypium hirsutum* (Tian et al., 2019); seed-cotton yield per plant is the fitness proxy. **(B)** and **(E)** Pr(*s <* 0): the proportion of homozygous introgressions that are deleterious;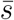: the mean effect of the homozygous interogressions; *r*: the estimator of eqs. 16-17 with a jackknife correction, eq. 35. **(C)** and **(F)**: “Prop. Dom” and “Prop. Overdom”: the proportion of QTL falling in the blue and red regions of the plot, and so supporting the dominance and overdominance theories;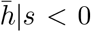 and 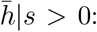 the typical dominance coefficient (eq. 9) for deleterious and beneficial introgressions. Bars are are 95% binomial confidence intervals for Pr(*s <* 0), and bootstrap CIs for all other statistics. ((C): means and standard deviations across replicate experiments from five years; (E)-(F): gray points show the individual years).

Figure 3A-C show data from Semel et al. (2006), comprising introgressions from the wild tomato species *Solanum pennellii* into the cultivated tomato plant *Solanum lycopersicum* – a species pair that shows F1 heterosis for some vigor- and fitness-related traits (Eshed and Zamir, 1995; Ghani et al., 2020). Figure 3D-F show comparable data from Tian et al. (2019), which the authors replicated in five years and two field sites. These data involve introgressions between the New World cotton species *Gossypium barbadense* and *Gossypium hirsutum*, which are also known to show F1 heterosis for many traits (Davis, 1979; Tian et al., 2019).

While Semel et al. (2006) and Tian et al. (2019) present data for multiple traits, we focus here on the traits with the largest number of reported QTL, namely seed weight per fruit (Figure 3A-C) and seed-cotton yield per plant (Figure 3D-F). While neither is a true component of fitness, each is classified as a reproductive or yield-related trait, and each is highly positively correlated with other such traits in these data (see Table 4 from Semel et al. 2006, and Table S1 from Tian et al. 2019). This is why, when comparing the data to the theory of eq. 3, it is reasonable to equate the measured traits with ln *w* (log fitness) rather than *z*_*j*_ (the idealized phenotypic traits with intermediate optima).

As Figure 3 shows, the results for the two systems show some strong similarities. Most importantly here, both data suggest a complex architecture of heterosis, with QTL appearing in multiple regions of the plot, and high variance in the selective effects. Moreover, the possible bimodality of the distributions (especially evident in Figure 3A) is plausibly attributed to difficulties in detecting QTL of small effect, which should cluster around the origin. As such, it is possible that both data sets represent a region of parameter space where the standard tests can be misled (Figure 2B and F).

With this in mind, let us first consider tests for coadaptation, which are summarized in Figure 3B and E. For both data sets, the proportion of QTL that are deleterious, Pr(*s <* 0), does not differ significantly from 1*/*2, implying no tendency for alleles to perform better in their homospecific background. While this test can be misleading (Figure 2D and H), stronger tests support the same conclusion. In particular, the mean effect of the homozygous introgressions,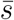, does not differ significantly from 0, and the estimator *r* (eq. 16 with a jackknife correction for bias; eq. 35) does not differ significantly from 1. As such, neither data set gives any evidence of substantial coadaptation between the homospecific alleles.

Now let us consider the evidence for overdominance, which is summarized in Figure 3C and F. When the variance in selective effects is high, the phenotypic model can generate overdominant QTL via transient overdominance (Figure 2B-C, F-G). However, this hypothesis does not provide a good quantitative fit to the data. First, when transiently overdominant QTL are present, they are always predicted to be rarer than QTL supporting the dominance theory (Figure 2C and G and Figure S1) – even when they are compound QTL (Figure 2B and G). Yet in the real data, overdominant QTL are in the majority (Figure 3C and F; with the sole exception being 1/5 years in the Tian et al., 2019 data). Similarly, transient overdominance can inflate the typical dominance coefficients (eqs. 10-11), but only within certain bounds. Even for compound QTL, when *D < n*, then eqs. 10-11 are bounded at *ĥ*|(*s >* 0) *<* 0.82 and *ĥ*|(*s <* 0) *>* 0.19, and yet the values for the real data are more extreme (Figure 3C and F; with only the deleterious introgressions in the Tian et al., 2019 data falling within the bounds). So while the phenotypic model can generate architectures with the same qualitative shape as those observed (Figure 2B and F, Figure 3A and C), it cannot account for such high levels of overdominance.

## Discussion

When F1 heterosis is fixable, hybridising populations can enjoy long-term fitness benefits. Heterosis should be fixable whenever the parental lines contain distinct sets of beneficial alleles that can be combined together as homozygotes (Mackay et al., 2020). But heterosis will not be fixable if there is true overdominance – an intrinsic advantage to heterozygosity – or coadaptation – a tendency for alleles to beneficial only in their original genetic background. Introgression lines can be used to test for both overdominance and coadaptation, and therefore for fixability.

We have studied a well-known model of phenotypic selection, which gives similar results to other such models (Martin, 2014; Cotto and Day, 2023). We have shown that this model can mislead the most straightforward tests of fixability – especially when there is a high variance in the selective effects, and variable fitness interactions between pairs of alleles (Figure 2). While the classical theories of heterosis have distinguished between true overdominance and associative-or pseudo-overdominance (Shull, 1908; East, 1908; Bruce, 1910; Keeble and Pellew, 1910; Jones, 1917; Collins, 1921; Crow, 1948), this model suggests a third possibility: transient overdominance (Sellis et al., 2011; Manna et al., 2011), where single mutations can be overdominant, but in a way that is strongly background dependent, such that the overdominance will come and go in recombinant hybrid genotypes. Transient overdominance, with or without coadaptation, can arise under the process posited by the classical dominance theory, i.e., the fixation of recessive deleterious mutations via genetic drift. It follows that a complex architecture of heterosis – even with very small QTL (Alseekh et al., 2013) – is not strong evidence against fixable heterosis, nor even against the process described by the classical dominance theory.

Our analyses also place bounds on the amount of apparent overdominance that can be explained in this way, and we have argued that the data of Semel et al. (2006) and Tian et al. (2019) exceed these bounds. As such, the alternative explanations for the data in *Solanum* and *Gossypium* remain true overdominance or pseudo-overdominance, and these alternatives remain very difficult to tell apart. Semel et al. (2006) and Tian et al. (2019) both argue for true overdominance, citing the high levels of heterosis, and the rareness of overdominant QTL in traits that are unrelated to fitness. These arguments are not, however, conclusive. The dominance theory can explain arbitrarily high levels of heterosis if mutations have reached fixation via drift (Crow, 1948; see also eq. 8), and the directional dominance that is necessary for pseudo-overdominance to appear, might be restricted to fitness traits (Mackay et al., 2020; Billiard et al., 2021). There is also suggestive evidence that heterosis is fixable in these species pairs. For example, high-fitness recombinant lines have been bred from *Solanum pennellii* and *Solanum lycopersicum*, (Christakis and Fasoulas, 2001; Gur and Zamir, 2015; Avdikos et al., 2021a,b); and *Gossypium barbadense* and *Gossypium hirsutum* show evidence of many successful introgression events (Wang et al., 2022), despite reports of strong fitness reduction for post-F1 hybrids (e.g. Jiang et al., 2000).

The studies of Semel et al. (2006) and Tian et al. (2019) were reanalyzed here because they present arguably the strongest evidence for multi-locus overdominance (Kaeppler, 2012). However, many other experimental approaches, and many other systems, have yielded evidence for overdominance (Yu et al., 1997; Birchler, 2003; Koltunow and Tucker, 2003; Lippman and Zamir, 2007; Jiang et al., 2017; Xie et al., 2022), and transient overdominance could, in principle, explain these data. It is notable, for example, that Xie et al. (2022) present evidence for the background dependence of overdominance in rice hybrids (*Oryza sativa japonica* and *O. s. indica*).

Finally, we note that phenotypic model presented here can also generate heterosis in other ways, including single locus overdominance (Manna et al., 2011) and negative epistasis between parental alleles (i.e. a reversal of the sign of *α*_2_ in eq. 15; see Hill, 1982), which is especially likely to arise under directional selection (Schneemann et al., 2020; De Sanctis et al., 2023). It can also generate heterosis that varies with environmental conditions (Schneemann et al., 2020, 2022). All of these have been observed in other systems (Moore, 1977; Eshed and Zamir, 1996; Wang et al., 1997; Krieger et al., 2010; Sang et al., 2022). As a result, the simple model used here might form the basis of a unifying theory of heterosis (Birchler, 2003; Kaeppler, 2012).

## Acknowledgments

JW is very grateful to Nicolas Bierne, Ali Duncan and Guillaume Martin for their help with the LabEx CeMEB (Centre Méditerranéen de l’Environnement et de la Biodiversité) exchange program, for which this work was developed. Both authors are also grateful to the authors of the tomato and cotton studies for making their data publicly available. HS acknowledges support from the Wellcome Trust program in Mathematical Genomics and Medicine (RG92770). JW also thanks the Popgroup56 audience who endured a presentation of this work with no data points on the slides.

## Appendix

In this Appendix we derive the theoretical results reported in the main text, describe the simulation procedures, report additional simulation results, and provide details of the curation of published data.

### Derivations

We begin with derivations that apply to the phenotypic model, regardless of how the parental lines diverged. First, let us denote as *z*^P1,*j*^ the difference between the optimal value of trait *j* (where *j* = 1, 2, …, *n*) and its actual value for the parental genotype P1. P1 fitness then follows directly from eq. 3

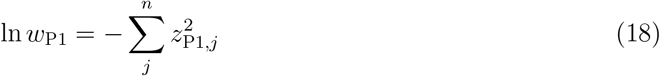

Note that this expression treats the P1 line as genetically homogeneous, and we will use this assumption throughout. An extension to within-line genetic variation follows from results in De Sanctis et al. (2023), but has no qualitative effect on the results reported here.

Now, let us consider introgressions of divergent alleles from donor line P2 to recipient line P1 (as depicted in Figure 1B). To calculate the selective effects of these introgressions, we need the additive and dominance effects of the *D* alleles that differentiate the lines. Denote these effects as *a*_*ij*_ and *d*_*ij*_, respectively, where *i* = 1, 2, …, *D* and *j* = 1, 2, …, *n*. Each heterozygous introgression will involve both effects, so if we consider the introgression of divergent allele *i* in heterozygous form, we have

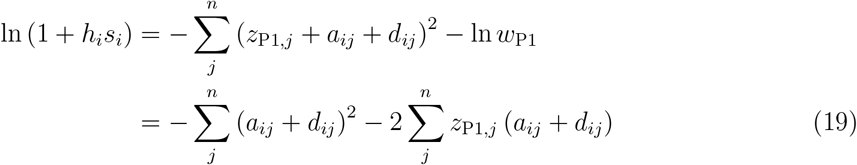

The equivalent homozygous introgression involves double the additive effect, and so we have

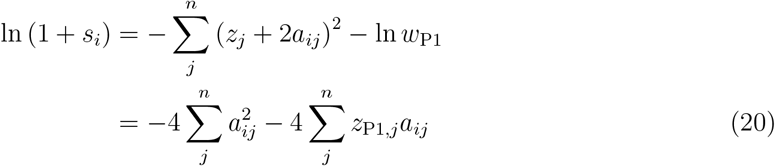

The proportion of QTL that support the overdominance and dominance theory (i.e. the proportion that fall in the red and blue regions of Figure 1C), can also be found from the bivariate distribution of heterozygous and homozygous effects. If we denote this distribution as *f* (*x, y*), where *x* = ln(1 + *h*_*i*_*s*_*i*_) and *y* = ln(1 + *s*_*i*_), then the proportions are given by the following integrals.

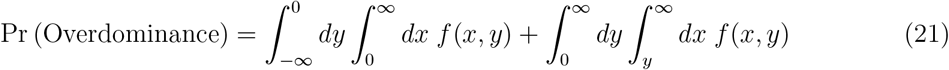

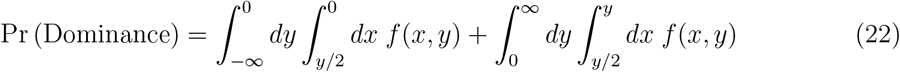

which are the quantities plotted in Figure 2C and 2G. The distribution *f* (*x, y*) itself is plotted in Figure 2A-B and 2E-F. For this purpose, we approximated *f* (*x, y*) with a bivariate shifted gamma distribution, with means and covariances appropriate for the model of divergence (see Table S1 below). We calculated the bivariate gamma distributions using the approach of Moran (1969), which maps the variables to a bivariate normal (Yue et al., 2001), and further assumed that the correlation coefficient of the gamma distributions was equal to that of the normals. The contours show the smallest region of the plot containing 95% of these probability densities. The proportion of QTL that are deleterious in homozygous form, as shown in Figure 2D and H, depends only on the homozygous effects, and is simply

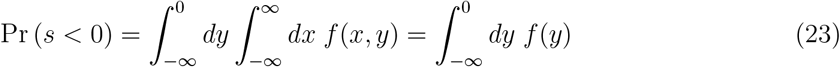

To calculate the level of heterosis, let us write the log fitnesses of the other parental line, P2, and the initial F1 hybrid (Figure 1A) by including the effects of all *D* alleles.

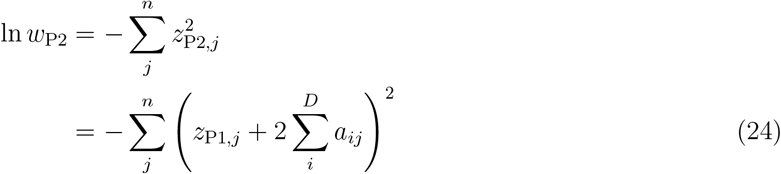

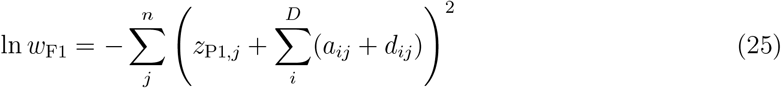

By combining eqs. 18, 24 and 25, our measure of F1 heterosis (eq. 8 with *π* = 1) is found to be:

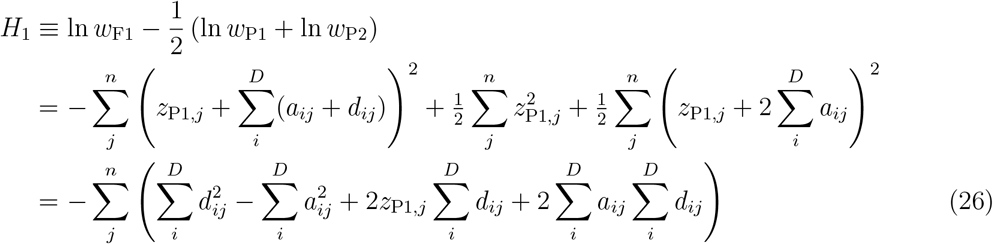

We can relate this work to the composite effects of Hill (1982) by using results from De Sanctis et al. (2023) (see also Schneemann et al., 2020), which show that the two-locus composite effects are given by

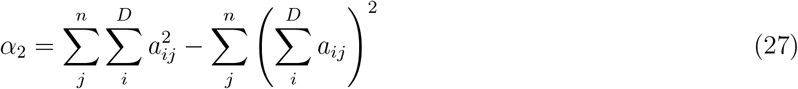

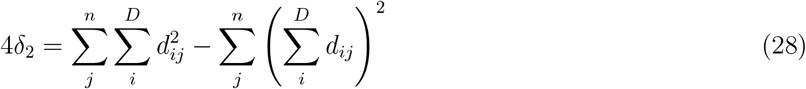

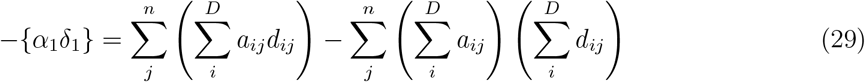

(note that {*α*_1_*δ*_1_} is the additive-by-dominance interaction, not the product of two single-locus terms). The single-locus dominance effect, *δ*_1_ = (*H*_1_ + *α*_2_)*/*2 cannot be written in such helpful form, but we can combine all of the results so far to note that:

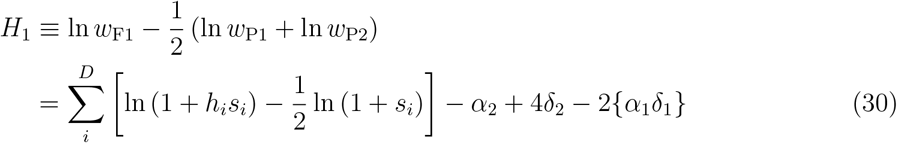

This is useful, because we will show below that the second-order composite effects involving dominance are zero on average (*E*(*δ*_2_) = *E*({*α*_1_*δ*_1_}) = 0) under both models of divergence used here (Schneemann et al., 2022; De Sanctis et al., 2023). Equation 15 of the main text then follows directly from eqs. 2 and 30 under the assumption that selective effects are small such that ln (1 + *s*) *≈ s* and ln (1 + *hs*) *≈ hs* – an approximation we use throughout in the main text. Finally, in the light of this approximation, we can write the estimator for the typical dominance coefficient (eq. 9) as

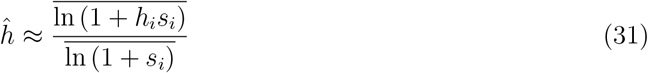

It follows, therefore, that eq. 16 can be written as

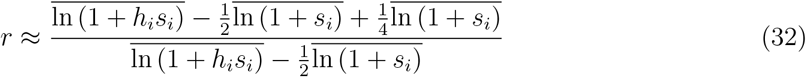

Now, it follows from eq. 30 that the expected value of the denominator of eq. 32 is

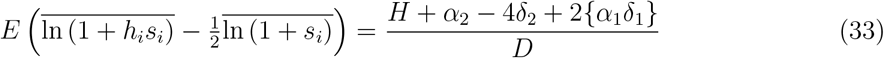

Next, it follows from eqs. 18 and 24 that 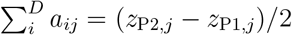. Factoring this result into eqs. 20 and 27, it follows that the extra term in the numerator has the following expected value.

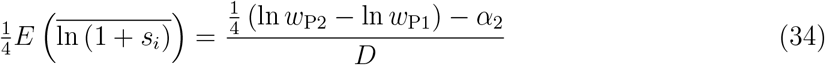

Equation 17 then follows from the additional assumptions (1) that the dominance effects have no tendency to point in a phenotypic direction (i.e., that *E*(*δ*_2_) = *E*(*α*_1_*δ*_1_) = 0); (2) that the two parental lines are similarly maladapted (i.e. that ln *w*_P2_ *−* ln *w*_P1_ *≈* 0). We note also that this ratio estimator (using the simple ratio of sample means) can be highly biased. There are various means of mitigating the bias (e.g., Blanchet et al., 2015), but in our data reanalysis (Figure 3), we use a simple jackknifing approach, as follows:

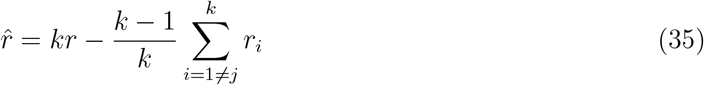

where *k* is the number of observations, and *r*_*i*_ is calculated omitting the *i*^th^ data point.

### Random mutation accumulation

Now let us consider results that require assumptions about the mode of divergence between the parental lines. To model random mutation accumulation, we simply assume that the additive and dominance effects of each fixed difference between the parental lines are independent normal variates. As such, we assume

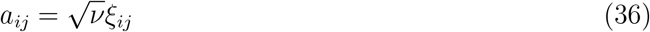

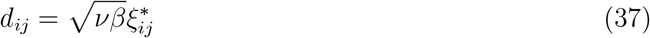

where *ξ*_*ij*_ and 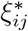 are independent standard normal variates. It follows immediately from standard normal theory that all three of the pairwise composite effects (eqs. 27-29) have expected values of zero under random mutation accumulation: *E*(*α*_2_) = *E*(*δ*_2_) = *E*({*α*_1_*δ*_1_}) = 0 (Schneemann et al., 2022; De Sanctis et al., 2023).

Now, let us assume that the ancestral genotype was phenotypically optimal, and that a proportion *p* of the *D* fixed differences went to fixation in the P1 lineage, while the remainder, (1 *− p*)*D*, went to fixation in the P2 lineage. It follows that the difference of the P1 phenotype from the optimum can be written as

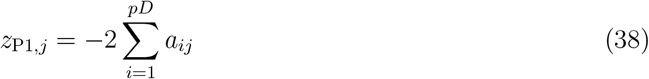

where we have ordered the additive effects, such that the first *pD* are the effects that fixed in the P1 lineage. From eqs. 2, 18 and 24-25, the expected heterosis is then:

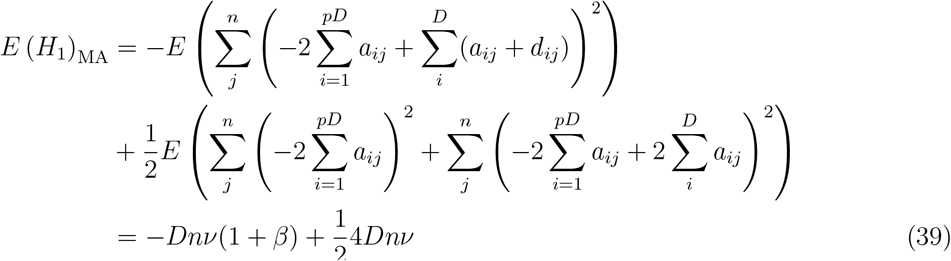

which, with eq. 4, yields eq. 8. Let us now consider a sample of introgressions where with probability *q*, the introgressed factor involves an allele that fixed in P1, so that the introgression from P2 is of an ancestral allele; 1 *− q* is therefore the probability that the allele fixed in P2, and is therefore an introgression of a derived allele (note that 1 *− q* is the quantity plotted on the x axis in Figure 2D). We can now use eqs. 19-20 with eqs. 36-38 and standard normal theory, to derive properties of the bivariate distribution of selective effects. For example, it follows from the assumptions above that

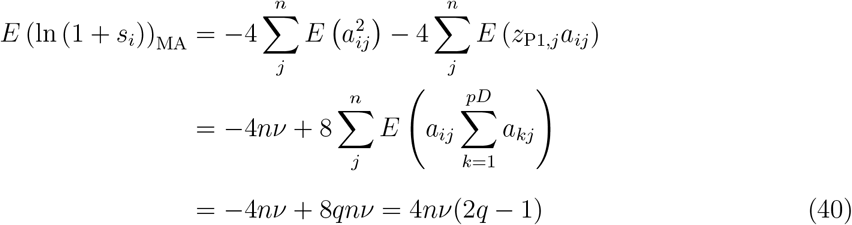

The variances and covariances follow by the same approach, and all of the relevant results are collected in Table S1A. These results were used to provide the means and covariances for the bivariate shifted gamma distributions shown in Figure 2A-B.

### Introgression of single QTL

The results also justify assertions in the main text. Let us first compare the introgressions of individual derived or ancestral alleles, by setting *q* = 0 or *q* = 1 in Table S1A. In this case, the typical selection coefficients are:

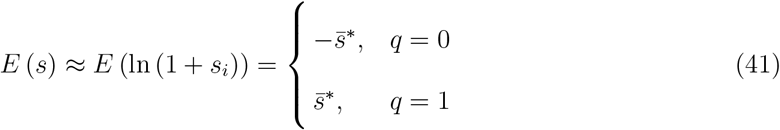

so under random mutation accumulation, derived alleles are deleterious on average, while ancestral alleles are beneficial. For the dominance coefficients, we find:

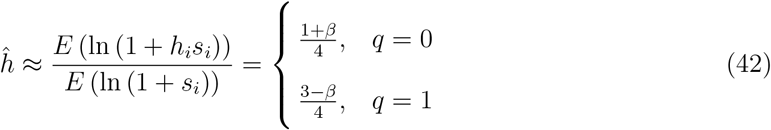

so that, if *β <* 1, derived alleles are recessive on average, and ancestral alleles dominant. If we consider the variances of the distributions for ancestral or derived alleles (such that *q*(1 *− q*) = 0), and assume that a reasonable number of differences have accrued in the recipient population (such that *pD ≫* 1), then we find that the standard deviations of the distributions scale with 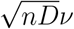, while the difference in means scales with *nν*. This implies that the distributions for derived and ancestral introgressions will be separate if *D ≪ n*, but overlapping if *D ≫ n*.

**Table S1:**
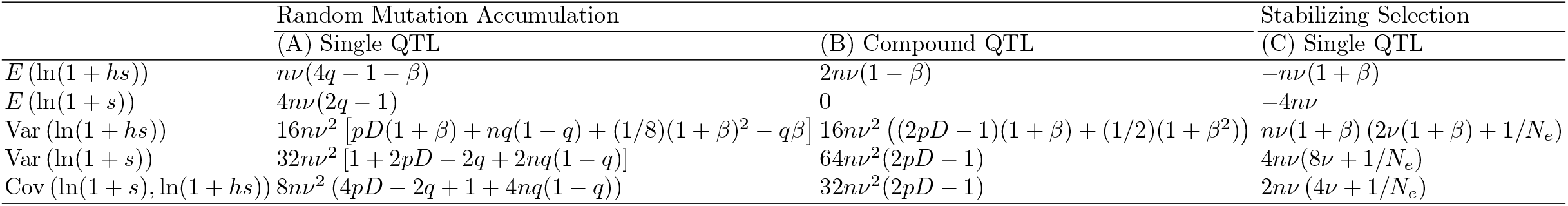
Means and covariances of introgression effects

In the first regime, where the distributions are separate (*D ≪ n*), then derived introgressions are almost certain to be deleterious and ancestral introgressions almost certain to be beneficial. It follows directly, that

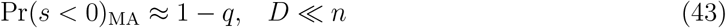

which is the 1:1 black line plotted in Figure 2D. It also follows that, when *D ≪ n*, eqs. 10-11 should match eq. 42, which shows that *c* = 1 in this case.

In the alternative regime, where *D ≫ n*, most of the analytical results are obtained by assuming that the bivariate distribution of heterozygous and homozygous effects (*f* (*x, y*) of eqs. 21-22) can be approximated by a bivariate normal, with means and variances as in Table S1A. For example, for the curves in Figure 2C, we start with eqs. 21-22 and results in Table S1A. We then (1) include both derived and ancestral introgressions in equal proportions, such that 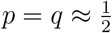, and (2) assume that *D ≫ n*, so that we can neglect terms in *n*^2^. In this case, the distribution *f* (*x, y*) has the following means: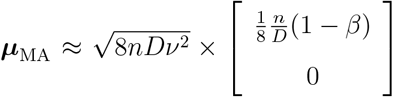, and (co)variances 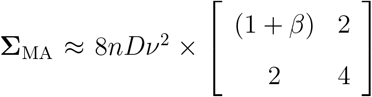. Now, with bivariate normality factors common to the squared means and covariances will have no effect on the result. This means that the proportions in eqs. 21-22 will depend solely on the two parameters found in the square brackets, namely *β* and the ratio *D/n*. Equations 21 and 22 can then be solved by numerical integration, and these results are plotted as the lighter solid lines in Figure 2C, corresponding to the simulations with *β* = 0.01 and *β* = 1*/*3. This approximation breaks down with phenotypic additivity, i.e. when *β* = 0, and so we use a different approach. First, we define the ratio *u*_*i*_

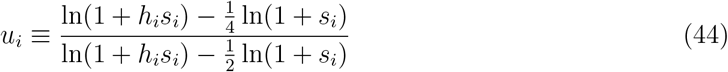

which is clearly related to the quantity *r* (see eq. 32). We also note that, under phenotypic additivity, QTL supporting the overdominance theory will show 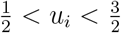, while all other QTL support the dominance theory. Now, using eqs. 36-38 with *β* = 0, we find

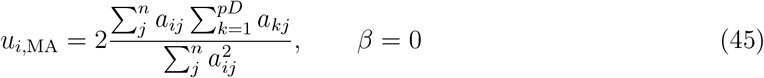

which, for *n* sufficiently large (say *n >* 3), is approximately normal with mean and variance given by

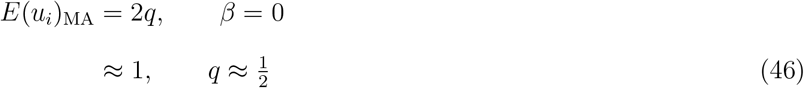

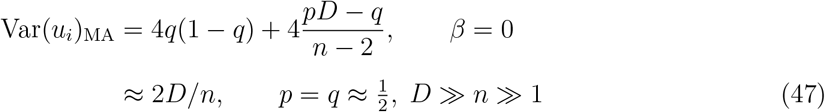

So, the probabilities in eqs. 21-22 can be written as

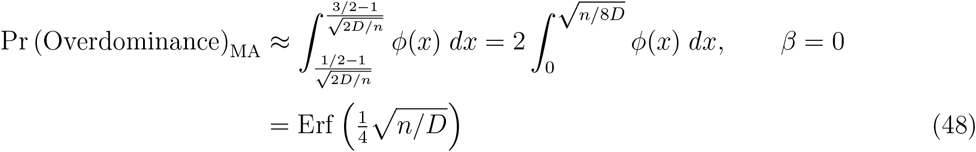

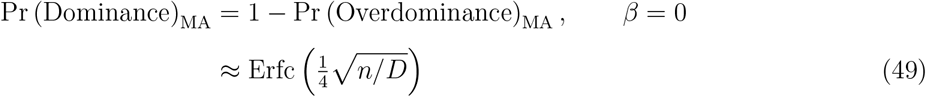

where *ϕ*(*x*) is the standard normal density, and Erf(*·*) and Erfc(*·*) are the error function and complementary error function (Abramowitz and Stegun, 1964). These results are plotted as the darker solid lines in Figure 2C, corresponding to the simulation data with *β* = 0.

The results in Figure 2D use a similar normal approximation for the distribution of homozygous effects (i.e. for *f* (*y*) of eq. 23). Using the mean and variance of ln(1+*s*) in Table S1A, and assuming that *p* = *q* (i.e. that introgressions are chosen at random from the divergent sites), then we find

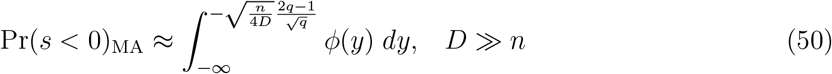

which is plotted against 1 *− q*, as the lighter curves in Figure 2D (and using the *D ≫ n* approximation for the simulations with *D* = *n*).

The same bivariate normal approximation is used to derive the values of *c* in eqs. 10-11, including results for the mean of a truncated bivariate normal (Rosenbaum, 1961). In particular, when *p* = *q ≈* 1*/*2, we have *E*(*y*) = 0 (Table S1A), and we find:

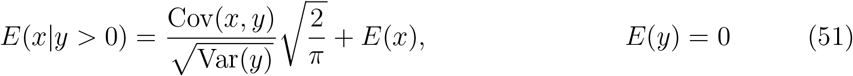

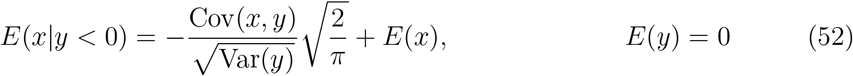

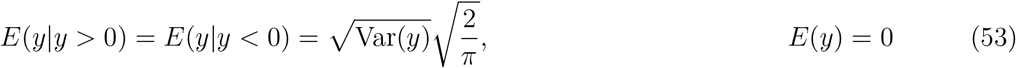

Equating *ĥ* with the ratio of these results (as in eq. 42), and using the moments in Table S1A, yields results that agree with eqs. 10-11, on the condition that 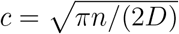.

### Compound QTL

All of the results above, which apply to single QTL, can be extended to compound QTL, containing exactly one ancestral and one derived alleles. For example, for an introgression containing derived allele *i* and ancestral allele *k*, we have

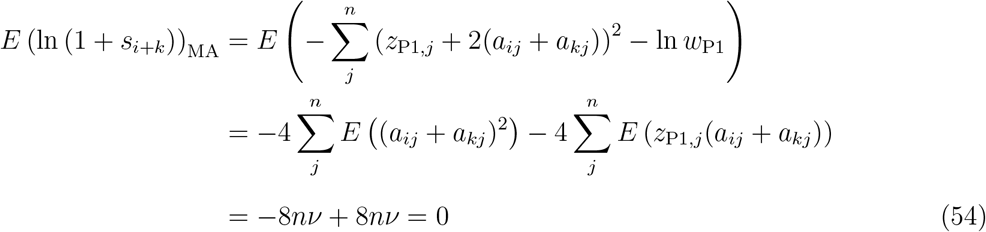

A complete set of results for such compound QTL are collected in Table S1B. These were used to generate the dashed lines and filled points in Figure 2A-B. Comparing results in Table S1A-B it is clear that when *D ≫ n*, results for compound QTL resemble those for single QTL, with 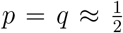, but with means and variances all approximately doubled. As such, if we assume of bivariate normality, then eq. 53, and the first term of eq. 52 will increase by a factor 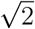, while the second term of eq. 52 will increase by a factor 2. If follows that eqs. 10-11 will also apply to compound QTL, but with 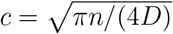.

#### Partially effective stabilizing selection

To model divergence under stabilizing selection, we must account for the fact that the fixed effects, which differentiate the parental populations, will not match the mutational input. However, predictions can be derived under the assumptions that the parental populations have reached a stochastic equilibrium under mutation, selection and drift, and that most of the genomic divergence has occurred during this phase, so that the shared ancestral phenotype has no influence on the results. In this case, the main effect of stabilizing selection can be captured by treating the parental phenotypes as random variables, and the additive effects as forming a tethered random walk between these phenotypes (Schneemann et al., 2020). In particular, we can write the additive effects as

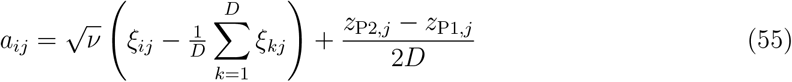

and the parental phenotypes as

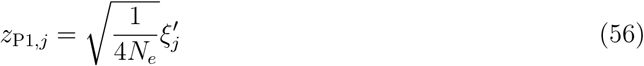

where 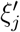 and *ξ*^*ij*^ are independent standard normal variates. It follows from eq. 56 that the expected parental fitnesses agree with the result of Barton (2016):

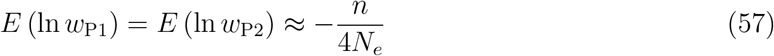

and that

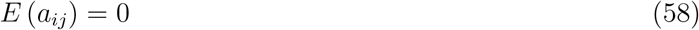

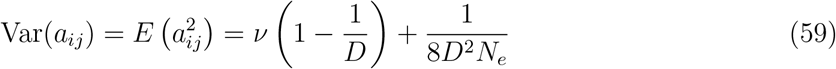

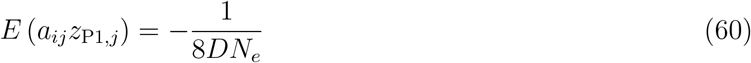

The expressions above apply only to the additive effects. Schneemann et al. (2022) and De Sanctis et al. (2023) show that, if the mutational correlations between the additive and dominance effects are not too strong, then the fixed dominance effects wander freely in phenotypic space, such that eq. 37 still holds under stabilizing selection. With the foregoing assumptions, and eqs. 27-29, the expected two-locus composite effects are found to be

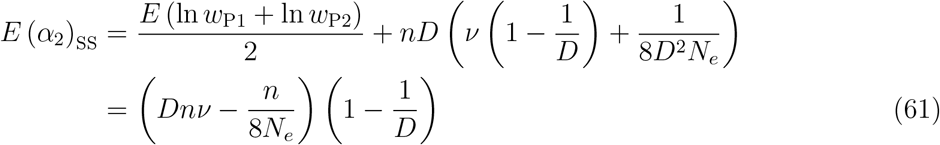

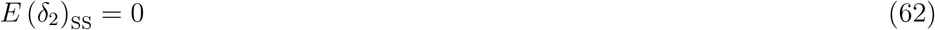

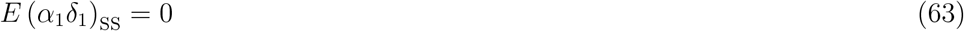

Equation 12 follows from eq. 61, after excluding terms *𝒪* (*D*^*−*1^). Equation 13 also follows, e.g., by using eq. 30. By using eq. 2, it can then be shown that eq. 5 still holds under stabilizing selection. By the same method, the homozygous and heterozygous effects of introgressions are found to be

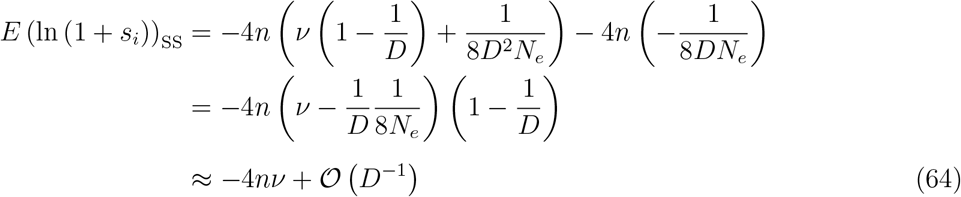

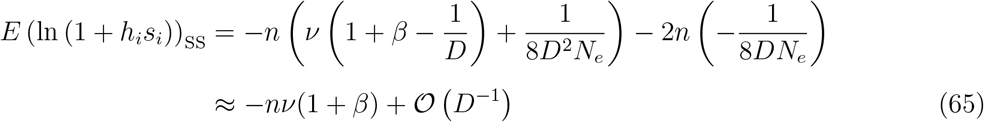

Expressions for the (co)variances of heterozygous and homozygous effects follow in the same way, and these results, to the same order of approximation (*𝒪* (*D*^*−*1^)), are collected in Table S1C. To produce Figure 2D-E, results in Table S1C were used with the bivariate shifted gamma approximation, described above. To calculate the curves in Figure 2F, we start with eqs. 21-22 and results in Table S1C, and then assume that *N*_*e*_*ν ≪* 1 su that we can neglect terms in *ν*^2^. In this case, *f* (*x, y*) has the following means: 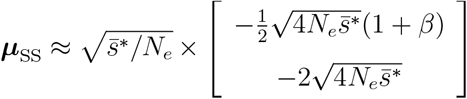, and (co)variances 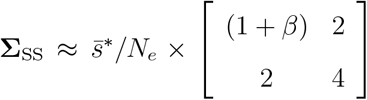, confirming that, when *N_e_ ν ≪* 1, the proportions of the QTL supporting the dominance and overdominance theories depend solely on *β* and 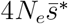 (Figure 2F). In the alternative limit, when *N*_*e*_*ν ≫* 1, we could use the same approach, but neglecting terms in 1*/N*_*e*_, yielding 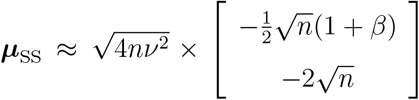, and (co)variances 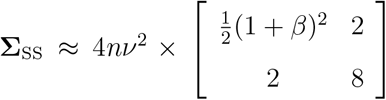. Note that in this regime, results depend solely on *β* and *n*. However, from eq. 4, the condition *N*_*e*_*ν ≫* 1 implies that 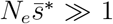, suggesting that this regime is less relevant to understanding complex architectures of heterosis, and so is not shown in Figure 2F. Figure 2F does, however, use a simpler approximation, which applies when *β* = 0, and is analogous to the approximation used under random mutation accumulation (eqs. 48-49). If we assume that *n* and *D* are sufficiently large, then once again, the ratio *u*_*i*_ (eq. 44) is roughly normally distributed, with approximate moments:

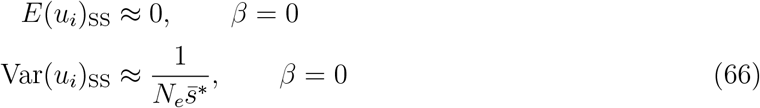

such that

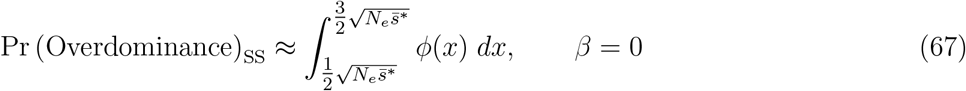

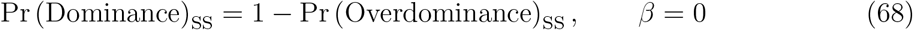

and these expressions are plotted as the darker lines in Figure 2F. Again, the proportion of homozygous introgressions that are deleterious (Figure 2H), also has a normal approximation, which uses the mean and variance for ln(1 + *s*) shown in Table S1C.

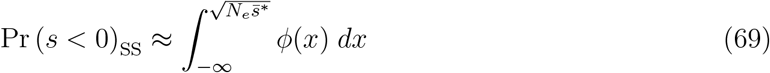

This result is plotted in Figure 2H. We note that this result is formally identical to the well-known result of Fisher (Fisher, 1930; Orr, 1998): for a given level of maladaptation (as determined here by the stabilizing selection), the probability that a mutation (or introgression) is deleterious, will increase with its size, from from 50% for very small mutations to 100% for very large mutations.

### Simulations

To generate the simulations of random mutation accumulation shown in Figures 2A-D and S2A, we generated *D* mutations, by drawing 2*Dn* normally distributed random numbers, using eqs. 36-37, and assigning a fraction *p* or 1 *− p*, to the two parental lines. We set *p* to *p* = 1*/*2 for all plots except Figure 2D, where it varied as shown on the x-axis.

These simulations and the analysis above all assume that the mutation effects on each trait are normally distributed. To understand the effects of violating this assumption, we carried out further simulations in which mutations sizes were exponentially distributed, and in which dominance effects were also non-normal. To do this, we replaced eqs. 36-37 with

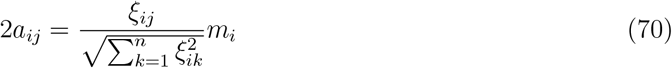

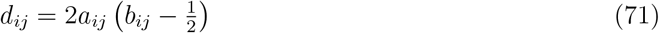

where *m*_*i*_ is an exponentially distributed random variable, with mean 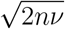, and *b*_*ij*_ is a beta-distributed random number with variance *β/*4. Equations 70-71 preserve the means and variances of eqs. 36-37 while relaxing the assumption of normality.

Results of simulations using eqs. 70-71 are reported in Figure S1A and Figure S2B. Results in S1A show that predictions for the architecture of heterosis (such as eq. 48-49) are only qualitatively robust to violating the normality assumption – i.e. the architectures are often complex, but the proportions in each region of the graph are poorly predicted (see also Fraïsse and Welch, 2019). By contrast, results in Figure S2B show that the use of *r* as an estimator (eq. 17) is robust to including non-normal mutations.

To simulate divergence under stabilizing selection, we used two approaches. The first, simpler approach, was to simulate the additive effects directly from eqs. 55 and 56, and the dominance effects from eq. 37. These simulation results are shown in Figures 2E-H and Figure S2C. We also extended this approach to non-normal effects by combining the bridge approach of eq. 55 with eqs. 70-71. Simulations generated in this way are shown in Figure S1B and Figure S2D. Again, as with random mutation accumulation, predictions for the architecture of heterosis are not robust to deviations from normality (Figure S1B), while predictions for *E*(*r*) remain robust (Figure S2D)

The simple bridge-based simulation approach allows us to control the size of fixed effects, and so cover the parameter space, but it does not model the action of natural selection explicitly. For this reason, we also compared our predictions to full individual-based simulations of stabilizing selection. The results of these simulations are reported in Figure S1C-D, and they all agree well with the simpler simulation approach.

The individual-based simulations followed the procedures reported by Schneemann et al. (2020) using custom scripts available at 10.5061/dryad.2bvq83bt9. In brief, we simulated pairs of Wright-Fisher populations of *N*_*e*_ diploid hermaphrodites, diverging from an optimal ancestral population through the joint action of mutation, selection and drift, in allopatry. Each discrete generation, parents were chosen with a probability proportional to their fitness (eq. 3). Each parent contributed a gamete with free recombination among loci. A Poisson-distributed number of mutations with mean 0.02*N* was then distributed randomly among all gametes (assuming infinite sites). The additive effects of these mutations on each trait were drawn from either a normal distribution (as in eq. 36; Figure S1C), or such that the total mutation magnitude was exponentially distributed (as in eq. 70; Figure S1D). The parameter *ν* was chosen such that the mean selection coefficient of homozygous mutation in an optimal background was 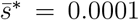. For all individual-based simulations, phenotypic dominance effects were generated with eq. 71. Simulations were run until one of the populations had fixed 10,000 substitutions, except for the simulations with *N*^*e*^ = 10 and *n* = 40 traits, which were slow to equilibrate, and so run until one population accrued 1 million substitutions. We ran simulations with three different population sizes (*N*^*e*^ = 10, 100 or 1000), and two different numbers of traits (*n* = 4 or 40), which combined with the three different levels of dominance (*β* = 0, 0.01, 1/3) and the two mutation models (eqs. 36 and 70). Fully crossed, this yielded 36 conditions in total.

After divergence, one population in each pair was randomly assigned as the donor for the simulated introgressions. Because system drift tends to prevent the fixation of larger-effect mutations, the simulations generated results only with smaller values of 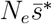. Accordingly, to extend results to higher values we also used the same simulation runs to generate compound introgressions by randomly bundling substitutions into groups of 10 or 100, and treating these as compound QTL. This method was used to generate the results with 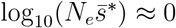 and 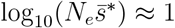 in Figure S1C-D. The bundling of fixed effects made their distributions closer to normal, explaining the differences between Figure S1B and S1D at higher 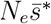 values.

**Figure S1:**
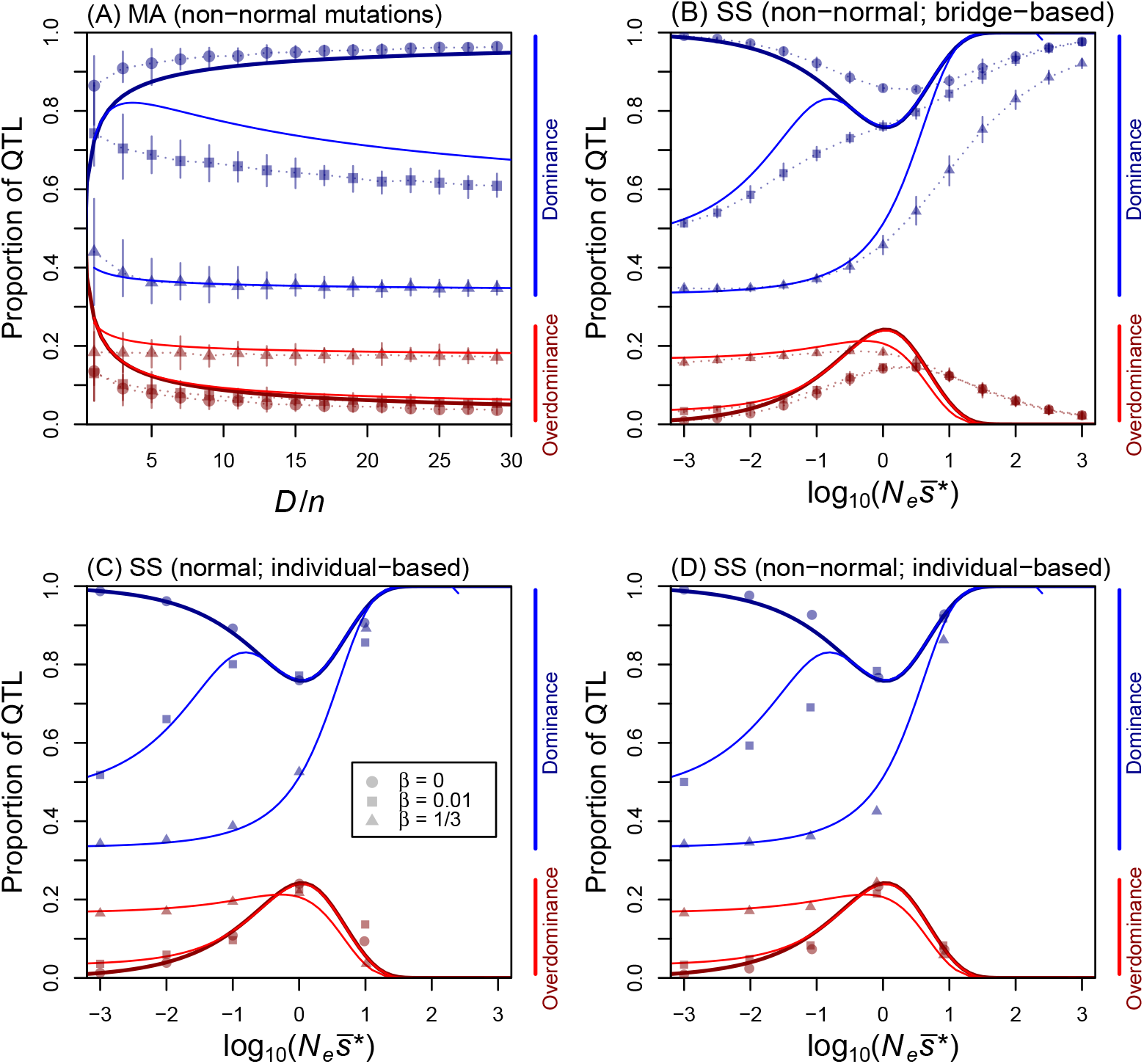
The architecture of heterosis when mutations are not normally distributed. **(A)** A replication of Figure 2C, but with the random mutations generated with eqs. 70-71 instead of eqs. 36-37, thereby violating the assumption that mutations have normally distributed effects. Curves show the predictions of eqs. 21-22 with values from Table S1A, and eqs. 48-49, which are shown to depend strongly on normality. **(B)** A replication of Figure 2G, but using non-normal substitution effects (eqs. 70-71) combined with the bridge-based simulation approach of eqs. 55-56 for the additive effects, to simulate the action of stabilizing selection. Curves show the predictions of eqs. 21-22 with values from Table S1C, and eqs. 67-68, which also depend strongly on normality. **(C)**-**(D)** Replications of Figure 2G, but using full individual-based simulations of divergence under stabilizing selection (see text for details). The additive effects of mutations were (C): multivariate normal (as in eq. 36), or (D) had exponentially-distributed sizes (as in eq. 70).

**Figure S2:**
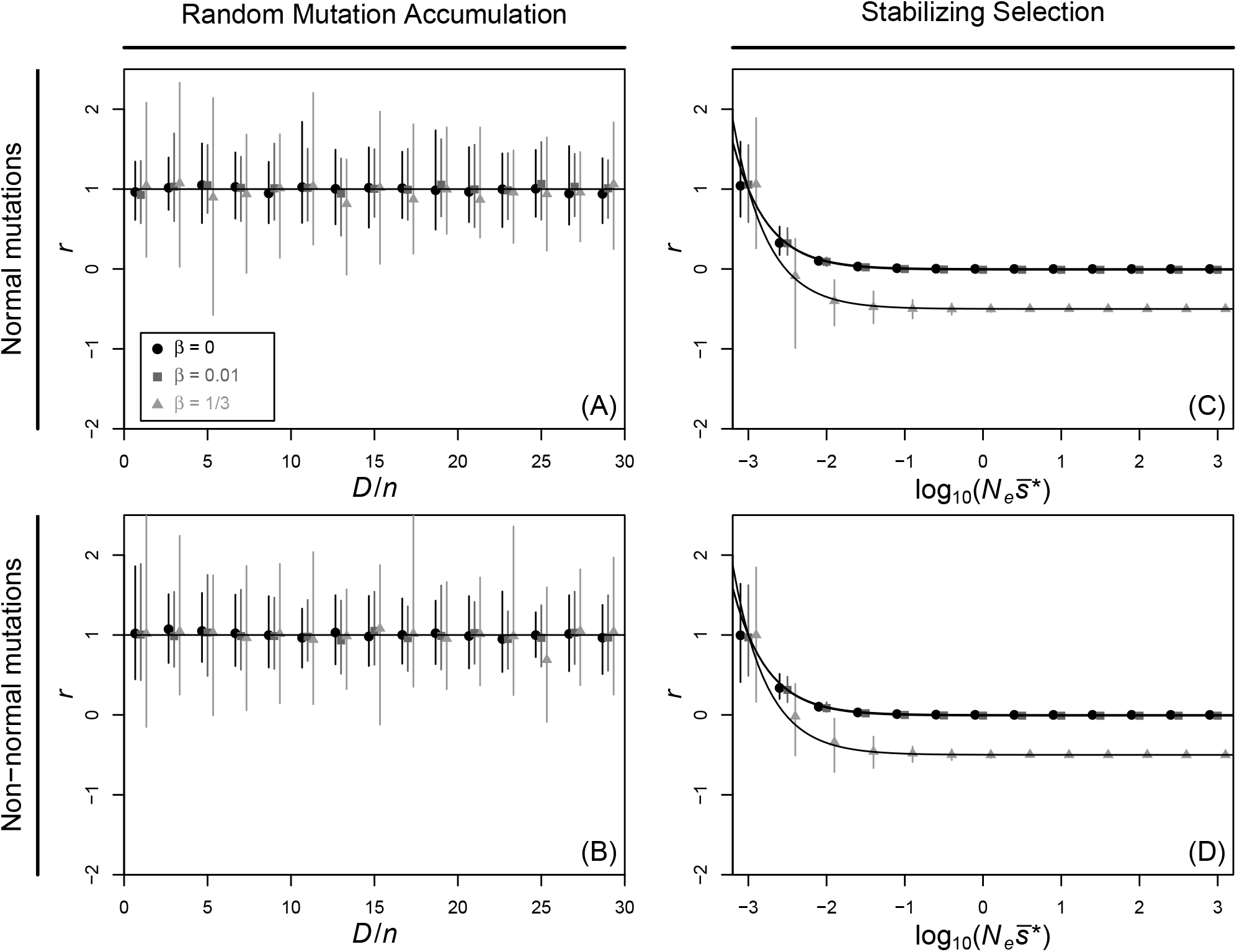
The scaled dominance coefficient can estimate the strength of coadaptation. Values of *r* (eqs. 9-16) plotted for various simulated data sets, with points and bars representing the mean and range across replicate simulations. **(A)-(B)** values are close to 1 when divergence takes place via random mutation accumulation. **(C)-(D)** values under stabilizing selection vary, as predicted from eq. 17 with eqs. 13 and 12 (curves), but 1 *> r ≥* 0 tends to hold for all simulation runs which generated both F1 heterosis (*H*_1_ *>* 0), and parental coadaptation (*α*_2_ *>* 0). The simulation runs shown are those found in (A): Figure 2C; (B): Figure S1A; (C): Figure 2G; (D): Figure S1B.

### Published data curation

This study was inspired by the observation of a complex architecture of heterosis for tomato introgression lines by Semel et al. (2006). We subsequently searched the literature for additional studies on heterosis that (i) reported data from an extensive set of (single segment) introgression lines, covering most of the donor genome, (ii) measured fitness or yield, and (iii) reported the fitness measurements from the introgression lines in both homozygous and heterozygous form. This search returned only the cotton study by Tian et al. (2019). To verify this, we searched web of science with the following search term: (ALL=(fitness) OR ALL=(yield)) AND ALL=(heterozygo*) AND ALL=(“introgression line*”) AND ALL=(heterosis).

The tomato introgression lines studied by Semel et al. (2006) were obtained from the Solanaceae Genomics Network (https://solgenomics.net/maps/pennellii_il/index.pl). These lines are comprised of inbred line M82 of the cultivated tomato variety *S. lycopersicum*, carrying single small partially overlapping chromosomal segments from the wild relative *S. pennelli*. The introgression lines from Tian et al. (2019) were created through several rounds of backcrossing Pima cotton (*G. barbadense*) to upland cotton (*G. hirsutum* line TM-1), followed by marker-assisted selection as described in Wang et al. (2012). Both studies compared the average trait values of individuals carrying a given introgression in homozygous or in heterozygous state, with that of the recipient line (i.e. M82 and TM-1, with averages taken across nine and twelve replicate individuals respectively). If any of these three values differed significantly, determined by a *t* -test with significance threshold 0.05 (or 0.01 for the comparisons to TM-1), the segment was recorded as harboring a QTL. The effects of introgressions that did not pass this significance threshold were not reported in these studies, and this might explain the apparent gap around the origin in Figure 3A and D. In the cotton study by Tian et al. (2019), this experiment was replicated at two locations (Dezhou and Jianpu Cotton Breeding Stations) and over five years (2010-2011 and 2013-2015).

